# Gene transcriptional profiling to assess Chinook salmon osmoregulatory resilience to salinity challenges

**DOI:** 10.64898/2026.09.30.755653

**Authors:** Shahinur S. Islam, Ken M. Jeffries, John W. Heath, Daniel D. Heath

## Abstract

Given unprecedented environmental change, Pacific salmonids are becoming increasingly threatened; however, genomic technologies provide an opportunity to assess Pacific salmonid resilience to on-going environmental change. Also, given increasing conservation and animal welfare concerns, non-lethal tissue sampling is emerging for assessing the physiological status of wild species. Relatively few studies, however, have compared the transcriptomic response of fish sampled non-lethally versus lethally for fish health and stress quantification. Here, we conducted a comparative gene transcriptional profiling study using 28 qPCR assays on the OpenArray® platform as an osmoregulatory stress transcriptional profiling chip. We sampled both non-lethal (gill biopsies, skin swabs, and water [eRNA]) and lethal (whole gill and liver) tissue samples from Chinook salmon (*Oncorhynchus tshawytscha*) smolts exposed for 0 h, 24 h, and 72 h to freshwater (0 PSU), brackish water (15 PSU) or seawater (30 PSU). We observed that most assays showed higher qPCR amplification success in both non-lethal gill biopsies and lethal gill tissues, compared to other tissues. We found more significant differences in transcript abundance in traditional lethal tissues compared to non-lethal tissues. With increasing duration of salinity exposure, gene transcript levels were typically upregulated in seawater samples compared to freshwater samples for both non-lethal and lethal tissues. The targeted genes for both non-lethal and lethal tissues were significantly and positively correlated across time points. Our findings support the use of non-lethal tissue sampling to improve Chinook salmon hatchery rearing, inform hatchery release timing, and facilitate future field application for the conservation of wild stocks.

## Introduction

Pacific salmonids are increasingly threatened by diverse environmental stressors. The impacts of climate change across the Pacific Ocean have driven widespread declines in Pacific salmon productivity and survival, affecting species throughout their freshwater and marine phases (Crozier et al., 2021; Metzger et al., 2026). Climate change-induced stressors such as increasing water temperature, decreased dissolved oxygen, changing coastal salinities, and droughts can impair Pacific salmon migration, elevate mortality rates, and increase vulnerability to disease (Evans et al., 2011; Bass et al., 2022; Akbarzadeh et al., 2024). These environmental stressors are further exacerbated by anthropogenic activities, posing an increasing threat to the persistence of salmon populations throughout their Pacific range (Miller et al., 2011; Peterman & Dorner, 2012). Many Pacific salmonid populations are in decline, with some already extirpated (Lehnert et al., 2019; Walters et al., 2019). Advanced genomic technologies can provide an opportunity to assess how Pacific salmonids will respond to rapid environmental change, identify molecular mechanisms underlying their stress responses, and evaluate the vulnerability of these species into the future (Healey, 2011; Crozier & Siegel, 2023).

Chinook salmon (*Oncorhynchus tshawytscha*) is an ecologically, culturally, and economically important species in British Columbia (BC), Canada. However, the sustainability of Chinook salmon fisheries are at risk due to substantial declines in the number of Chinook salmon throughout BC. Many factors contribute to these declines, but the timing and pace of smoltification have likely played a pivotal role (Handeland et al., 2004). During smoltification, juvenile anadromous Chinook salmon undergo extensive physiological changes for the transition from fresh water to sea water (Houde et al., 2019). These changes include modification of their osmoregulatory mechanisms, adjustments in metabolism, behaviour, and immunity, and shifts in endocrine signaling (McCormick et al., 1998; Quinn et al., 2005). Ill-timed seawater entry or inadequate smoltification can cause over 40% mortality and stunted growth (Stien et al., 2013). To increase wild Chinook salmon stocks and augment capture fisheries, aquaculture and hatchery breeding programs are used (Riddell et al., 2024). However, aquaculture- and hatchery-produced juvenile Chinook salmon generally have a lower seawater tolerance during their migration from fresh water to sea water, suggesting that the smoltification window may be altered in a hatchery environment (Shrimpton et al., 1994). While much is known about the physiological mechanisms of smoltification, less is understood about the molecular mechanisms (but see Houde et al., 2019; Akbarzadeh et al., 2020) underlying osmoregulatory resilience to environmental change in Chinook salmon.

The development of gene transcriptomic technologies has advanced the field of physiological genomics over the past decades, enabling assessment of gene expression that mediates species’ responses to aquaculture-relevant and environmental stressors (Schulte, 2004; Yue & Wang, 2017; Robledo et al., 2018). Transcriptional profiling quantifies gene transcription across multiple functionally related genes to assess targeted gene responses to specific stimuli (Semeniuk et al., 2022). As such, this is a powerful approach to assess physiological status of captive fish and the adaptive capacity of wild fishes (Connon et al., 2018). Comparative transcriptome profiling of targeted genes involved in osmoregulation can further identify the molecular signatures indicative of stress during seawater acclimation (Lee et al., 2020). Moreover, the application of advanced transcriptomic tools allows aquaculture and hatchery managers to detect early signs of stress, which is critical for aquaculture and fish welfare in general (Eissa & Wang, 2014). Furthermore, transcriptomics can provide important insights into the molecular genetic signatures underlying physiological plasticity (Wellband & Heath, 2017; Oomen & Hutchings, 2022; Banousse et al., 2025).

The application of transcriptomic approaches in fish health and conservation physiology has grown significantly over the last few years; however, there is still a lack of examples of the uptake this technology by aquaculturists and fisheries managers (Cooke et al., 2012). A key criterion for managers to embrace genomic technologies is to show how they can contribute to effectively managing resources experiencing anthropogenic activity impacts for sustainable exploitation (Friedman et al., 2022; Semeniuk et al., 2022). For example, for decades, researchers have been using peripheral tissues, such as liver, kidney, and heart to obtain mRNA (the RNA that is translated into protein yet makes up a small proportion of the total RNA) to assess cell, tissue and organism function and health. However, using tissues that require lethal sampling raises animal welfare concerns (e.g., for better husbandry practices) as well as conservation and management concerns (e.g., for threatened and endangered species). Researchers have thus started to use non-lethal tissue samples for transcriptomic studies (Jeffries et al., 2014, 2021). One major benefit of using non-lethal tissue sampling is the ability to integrate physiological genomics data with fitness-relevant ecological data such as reproduction, migration, growth, and survival (Thorstensen et al., 2022). Moreover, another advantage of using non-lethal tissue sampling is the ability to repeatedly sample the same individuals in chronic stress physiology studies (Teffer et al., 2019). For example, gill biopsies, one of the non-lethal tissue sampling methods used for fishes, have been used for decades (see Jeffries et al., 2021). An exciting emerging non-lethal sampling methodology is water or environmental RNA (eRNA) sampling. As a single-stranded molecule, RNA is less stable and degrades more quickly compared to double-stranded DNA (e.g., environmental DNA [eDNA]) making eRNA analysis more technically demanding; however, eRNA may better reflect the spatial or temporal signal of living organisms, leading to precise quantification relative to eDNA (Yates et al., 2021). What is less clear, however, is whether the transcriptomic profiles of non-lethal versus lethal tissues are similar for quantifying acute stress in fish.

Here, we describe using both non-lethal (gill biopsies, skin swabs, and water for eRNA) and lethal (whole gill and liver) samples associated with Chinook salmon smolts subjected to 0, 24 and 72 h acute salinity challenges (freshwater [0 PSU], brackish water [15 PSU], and seawater [30 PSU]). We conducted a comparative gene transcriptional profiling study using 28 TaqMan qPCR assays on the OpenArray® platform as an osmoregulatory stress transcriptional profiling chip. The main objective of this study was to quantify the gene transcriptional profiling to assess the resilience of Chinook salmon smolts under environmental change. Our secondary goal was to assess the transcriptomic responses of non-lethal versus lethal tissue samples, with the hypothesis that they would be equivalent under salinity challenge. This study will improve the assessment of the physiological status of Chinook salmon in aquaculture and hatcheries and foster better husbandry by enabling the screening of aquaculture and hatchery stocks for important production and performance traits, thereby enhancing production efficiency, profitability, and sustainability.

## Materials and Methods

### Study stock and sampling

This study was conducted between November 11 and November 13, 2022, at the Yellow Island Aquaculture Ltd. facility (Heriot Bay, BC, Canada). The Chinook salmon stock originated from the Fisheries and Oceans Canada hatcheries at Robertson Creek (BC, Canada) and Big Qualicum on Vancouver Island (BC, Canada) in the mid 1980s. The fish used for this study were from the 2021-year class and were being reared in freshwater raceways. Fish that reached the smolt stage were selected for this experiment (assessed by visual identification of loss of parr marks and intense silver colour). Water quality parameters in the experimental tanks were maintained at an average dissolved oxygen (DO) saturation of 99.89%, a DO concentration of 12.80 ± 0.03 mg/L, and a water temperature of 4.98 ±0.17 °C. Chinook salmon were placed in each of 12 experimental tanks (n = 40 fish tank^-1^; water capacity of each tank was approximately 200L) across three treatments (i.e., four tank replicates): freshwater (hereafter “FW”), brackish water (hereafter “BW”), and seawater (hereafter “SW”) (Figure 1). Careful consideration was given to minimizing stress associated with transportation and handling. The salinity for FW (controlled), BW, and SW was set at 0 PSU, 15 PSU, and 30 PSU, respectively. We collected both non-lethal tissue (gill biopsies, skin swabs, and water) samples and traditional lethal tissue (gill and liver) samples at three time points: 0 h, 24 h, and 72 h after salinity exposure. Before sampling, fish were euthanized with an overdose of clove oil (Aromatics Canada Inc; 0.4 mL L^-1^ clove oil in aerated hatchery water). Fish fork length and weight was measured prior to tissue sampling (see Supplementary Table S1). Lethal and non-lethal tissue samples were then collected at the time points from eight fish in each of the 12 experimental tanks. Briefly, for gill biopsy samples, fine-tipped scissors were used to excise only the distal tips of gill filaments, whereas whole-gill samples included gill filaments, gill raker, and gill arch. Liver tissue was collected using standard fish dissection procedures. For water/eRNA samples, we collected 500 mL of water in a sterile Nalgene bottle from each experimental tank at each of the three sampling time points. We also collected an additional 500 mL of water from each tank as a second filtration replicate. Additionally, we used 500 mL of ddH_2_O as a filtration control for each tank. We then filtered the water samples immediately using 47 mm diam Whatman^®^ glass microfiber filter (Millipore Sigma, Sigma-Aldrich, ON, CA), with a peristaltic pump (Geotech Environmental Equipment Inc, Denver, CO, USA). We cut the filters in half with scissors sterilized with a bleach solution and stored each half of the filter separately in 1.5 mL microcentrifuge tubes. For skin swab samples, the lateral surface of each fish was swabbed from the tip of the mouth to the base of the caudal fin using a sterile Fisherbrand™ swab (Fisher Scientific, ON, CA). Following sampling, the swab tip was removed and retained for subsequent molecular analysis. Collected lethal and non-lethal tissue samples were preserved immediately in a dry shipper with liquid nitrogen and later placed in a - 80 °C freezer until extractions. All fish were treated following the guidelines provided by the Canadian Council on Animal Care during holding and experimentation, and approval was granted by the University of Windsor Animal Utilization Project Proposal (AUPP#23-06).

**Figure 1:**
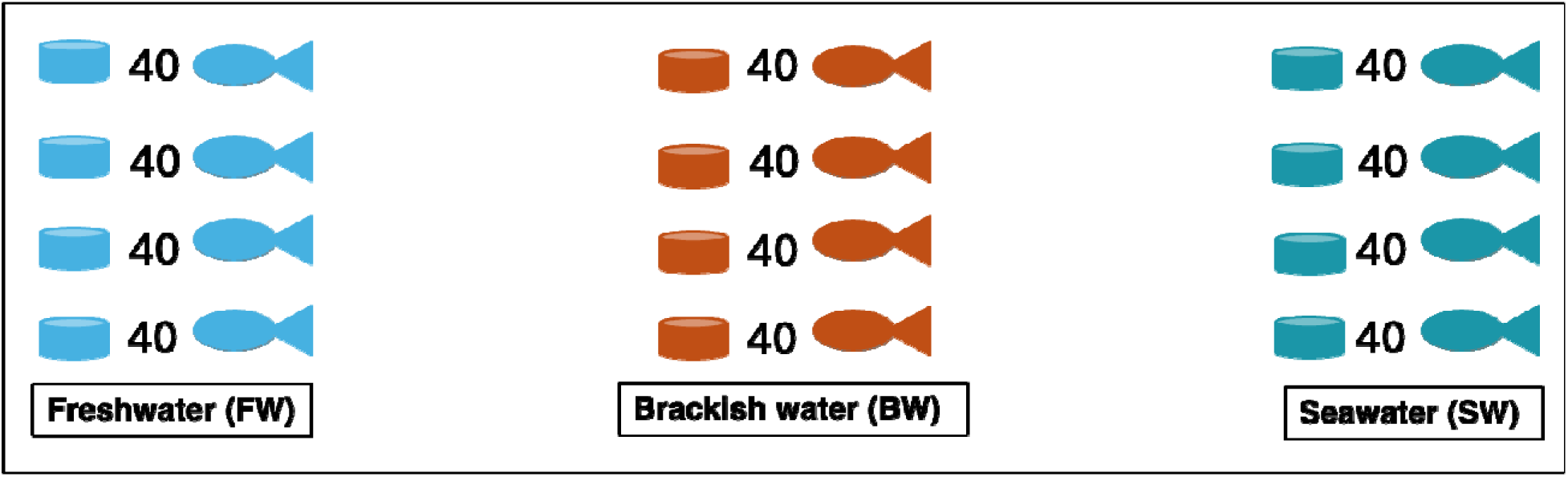
A schematic diagram of the experimental design. Forty Chinook salmon (*Oncorhynchus tshawytscha*) (smolt-stage) were placed in each of 12 experimental tanks across three treatments: FW (controlled), BW, and SW.

### RNA extraction and cDNA synthesis

RNA was extracted from the gill biopsies, skin swabs, water, gill, and liver tissues from a total of 360 samples (n = 3 treatments x 3 time-points x 4 tank replicates x 2 fish per tank x 5 tissues). Gill biopsies, gill, and liver tissues were extracted using TRIzol (Invitrogen/Life Technologies) with glass mill beads (1 mm, BioSpec Products) using a Beadbeater (BioSpec), following the manufacturer’s protocol. For extraction of skin swab samples, 700 µL of Proteinase K digestion buffer and 17.5 µL of Proteinase K were added to submerge the swab in a 2 mL tube and incubated at 55 °C for 1 h. After incubation, the liquid and swab were placed into an investigator basket (QIAGEN Investigator Lyse & Spin Basket Kit) and centrifuged at 17,000 g for 3 min at room temperature. The flow-through was processed using TRIzol, following the manufacturer’s protocol. For extraction of water samples, one of the two filter halves was used. Briefly, 700 µL of 1X DNA/RNA Shield^TM^ (Zymo Research), 80 µL of PK Digestion Buffer, and 20 µL of Proteinase K were added to completely submerge each half filter in a 2 mL screwcap tube and incubated at 55 °C for 1 h. After incubation, the liquid and filter were placed into a QIAshredder spin column (QIAGEN) and spun down at 17,000 g for 3 min at room temperature. The flow-through was then processed using TRIzol, following the manufacturer’s protocol. All extracted RNA was stored at -80 ^°^C. Total RNA integrity was checked using gel electrophoresis, and RNA purity was evaluated using A260/280 and A260/230 via NanoQuant for the SPARK plate reader (TECAN). Only samples with high purity (A260/280 > 2.0, A260/230 > 1.8) and integrity (clear 18S and 28S RNA bands) were used for cDNA synthesis. A total of 2 μg of total RNA was synthesized into c DNA using a High-Capacity cDNA Reverse Transcription (RT) kit (Applied Biosystems, Burlington, ON, Canada). The cDNA was stored at -80 ^°^C until further use.

### Candidate gene selection, chip design and optimization

This study used previously designed and optimized 28 TaqMan^TM^ qPCR assays for the OpenArray® platform as an osmoregulatory stress transcriptional profiling chip (Tugui et al., 2025; Islam et al., 2026). The chip included twelve genes involved in osmoregulation: cystic fibrosis transmembrane conductance regulator (*cftr*), sodium-potassium ATPase alpha subunit (*atp1a1*), sodium-potassium ATPase subunit beta-1 (*atp1b1*), solute carrier family 2-member 11a (*slc2a11a*), solute carrier family 9 member A1B (*slc9a1b*), solute carrier family 9 member 2 (*slc9a2*), solute carrier family 9 member A3 (*slc9a3*), solute carrier family 12 member 1 (*slc12a1*), solute carrier family 12 member 2 (*slc12a2*), ATPase H+ transporting lysosomal V1 subunit B, member a (*atp6v1ba*), Rh family B glycoprotein (*rhbg*), and Rh type C glycoprotein (*rhcg1*). Five gènes involved in immune function were included: interleukin 1β (*il-1*β), interferon γ (*ifn*γ), tumor necrosis factor α (*tnf-*α), mx protein (*mx*), and tapasin (*tapbp*). Four genes related to growth and metabolism were included: growth hormone receptor *(ghr)*, insulin-like growth factor 1 (*igf1*), insulin-like growth factor binding protein 1 (*igfbp1)*, and insulin-like growth factor binding protein 2 (*igfbp2)*. The chip also included four key genes involved in stress response: glucocorticoid receptor 1 (*gr1)*, glucocorticoid receptor 2 (*gr2*), 11β-hydroxysteroid dehydrogenase 2 (*hsd11b2*), and heat shock protein 90 (constitutive) (*hsp90ba*). Finally, three reference genes were included: glucose-6-phosphate 1-dehydrogenase (*g6pd*), elongation factor 1 alpha (*ef1a*), and ribosomal protein L13a (*rpl13a*). Additional information for each assay, including primer and probe sequences, amplicon size, accession numbers, and salmonid species used in the final alignments for primer and probe design, is provided in Supplementary Table S2.

More specifically, the Atlantic salmon ortholog (or ortholog from other salmonid) was searched for using BLASTn (NCBI) for sequence identification, alignments, and primer/probe design using Geneious Prime software (version 7.1.5; Kearse et al., 2012) (see Tugui et al., 2025; Islam et al., 2026 for details on primer and probe design). All primers and Taqman^TM^ probes for the 25 candidate and three reference gene assays were developed using Primer Express Software v3.0 (Thermo Fisher Scientific) for use on the OpenArray® chip platform. This chip was used to quantify transcription on a QuantStudio 12K Flex Real Time PCR System following the manufacturer’s protocol. Each chip could run 48 samples with qPCR duplicates (a total of 96 reactions) for 28 qPCR assays. For 360 cDNA samples, a total of 8 chips were used.

### Quantitative PCR (qPCR) assessment

A 5 µL mixture was prepared for each cDNA sample, which contained 2.5 µL TaqMan® OpenArray® Real-Time PCR Master Mix (Applied Biosystems, Canada), 1.3 µL ddH_2_O, and 1.2 µL cDNA. The 5 µL mixtures were prepared in 384-well plates and loaded into OpenArray® chips using the OpenArray® AccuFill System (Applied Biosystems) to minimize the inter-assay variation. The samples were randomly distributed, and all chips were normalized to the internal reference genes using ExpressionSuite Software v.1.0.3 (Applied Biosystems, Burlington, ON, Canada). The stability of the three reference genes was evaluated using RefFinder (Xie et al. 2023), which integrates four commonly used algorithms—geNorm, Normfinder, BestKeeper, and Delta C_T_—to compare and rank candidate reference genes. The gene *ef1a* (1.19) was the most stable assay, followed by *g6pd* (1.86), and *rpl13a* (2.28). We calculated PCR efficiencies for each assay for all five tissues (gill biopsies, skin swabs, water, gill, and liver) using LinRegPCR (Ramakers et al. 2003) (see Supplementary Table S3). The performance of qPCR assays for cDNA from each tissue is provided in Table 1. For estimating PCR efficiencies, we exported the raw amplification data from the QuantStudio 12K Flex Real-Time PCR system and separated the imported data by target gene to calculate an individual Window-of-Linearity for each target gene across all samples. For each assay, the mean C_T_ value (fractional PCR cycle number at which fluorescence crosses the quantification threshold) was used. We then estimated the starting cDNA concentration (N_0_ value) in each sample using the efficiency values (Tuomi et al., 2010) using the following equation:

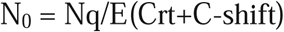

**Table 1:**
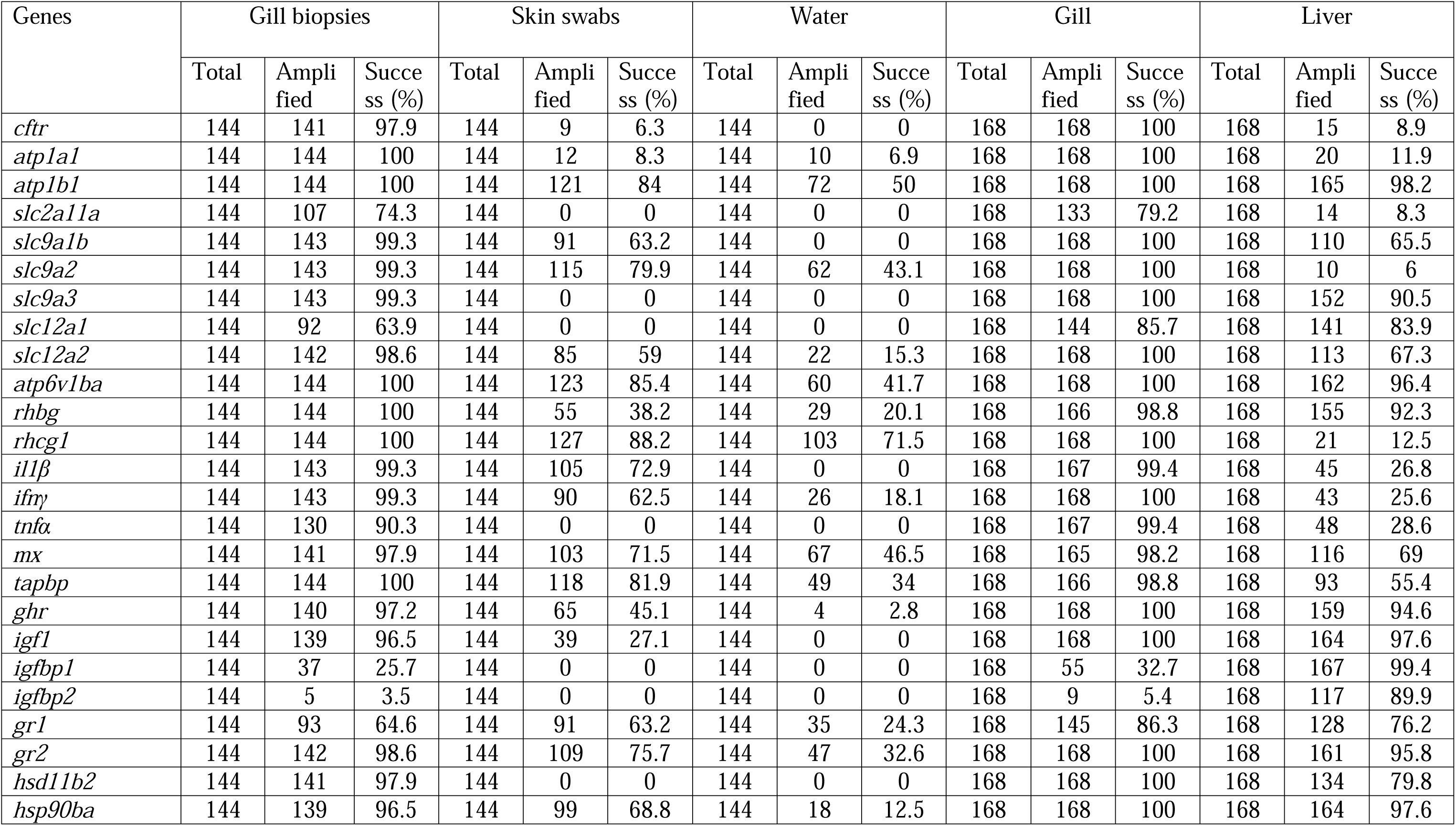
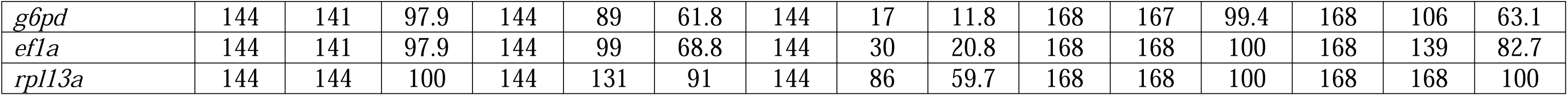
The performances of qPCR assays for cDNA from gill biopsies, skin swabs, water, gill, and liver tissues. The performance of qPCR assays was based on the success (%) of amplified samples for all treatment-by-time combination.

Where Nq represents the quantification threshold, set at 10^6^, E is the PCR efficiency, Crt is the threshold value of the gene, and C-shift corrects for different primer efficiencies. The C-shift value was calculated using the following formula:

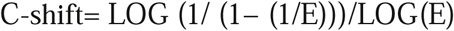

For relative transcription level, we then used the N_0_ value, divided by the geometric mean of the N_0_ of reference genes, to generate the ΔC_T_. We calculated ΔΔC_T_ values by averaging the ΔC_T_ values of the controlled FW fish and subtracting the ΔC_T_ of each individually BW and SW fish by the average of ΔC_T_ for that FW fish, accounting for PCR efficiency differences in our primers. We calculated ΔΔC_T_ only for those genes that showed a significant treatment/time-point and treatment x time-point effects.

### Statistical Analyses

All statistical analyses were performed in R version 4.5.0 (R Core Team, 2025). We performed linear mixed-effects (LME) models with the ‘lme4’ package (Bates et al., 2015) to detect transcript abundance differences across tissues (gill biopsies, skin swabs, water, gill, and liver; Table 2). Treatment and time point were included as fixed effects, and tank ID was included as a random covariate in the model. The final model was selected from the full model with the ‘lmerTest’ package, which allowed for automatic model selection (Kuznetsova et al., 2017). The ‘step’ function was used to perform backward selection. Non-significant random intercept was removed first, followed by non-significant interaction term, and then non-significant fixed effects. When an interaction term was significant, all fixed effects were included in the final model regardless of their significance level. Significance values for the fixed effects and interaction term were obtained using an F-test based on Satterwaite’s approximation. The final model was confirmed using plots of the model residuals, and the normality of the fitted model residuals was confirmed visually using histograms. In the final model, genes that showed significant differences in transcript abundances (ΔΔC_T_) were presented using graphical presentations for lethal gill and liver tissues (see figures 2-3). Tukey-adjusted multiple comparisons posthoc tests were performed for pairwise comparisons within each time point across tissues. The qPCR datasets (ΔC_T_) for the different tissues at three different time points were analyzed via principal component analysis (PCA) using the Factoextra R packages. This was completed using the ‘missMDA’ package with the ‘estim_ncpPCA’ function, which handles missing values in multivariate data analysis (Josse & Husson, 2016). The correlation coefficient (R) between the transcript abundance in lethal tissues and non-lethal tissues were also estimated using either Pearson (parametric) or Spearman (non-parametric) test and visually represented (see Figure 4).

**Figure 2:**
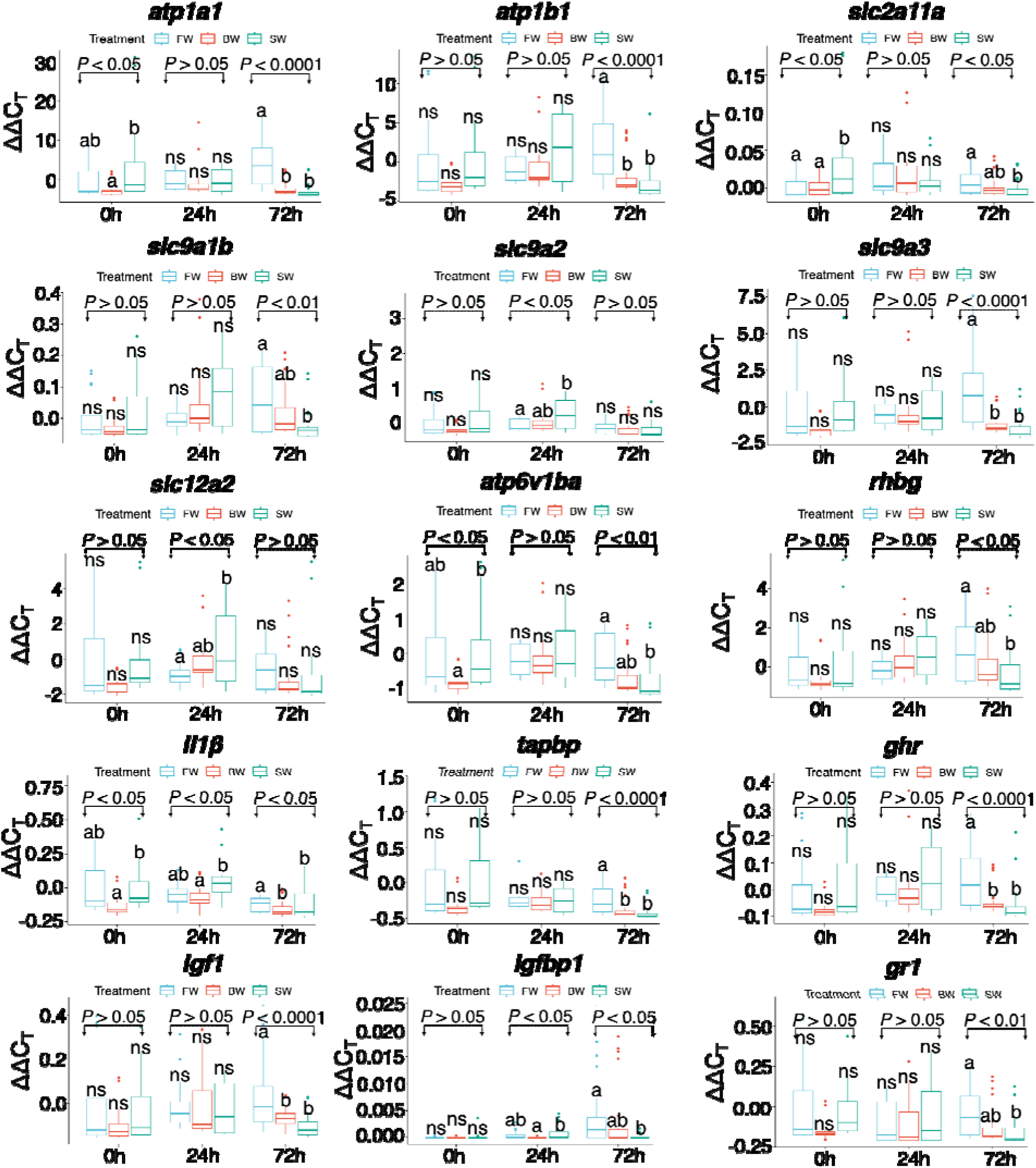
Box plots represent 15 genes that showed significant differences in transcript abundances in gill tissue based on mixed-effects models. The bold line represents the median, the boxes show the 25^th^ and 75^th^ percentiles, the whiskers indicate the 95% confidence interval, and the dots are outliers. Different letters denote significant differences in mean ΔΔC_T_ values among treatments.

**Figure 3:**
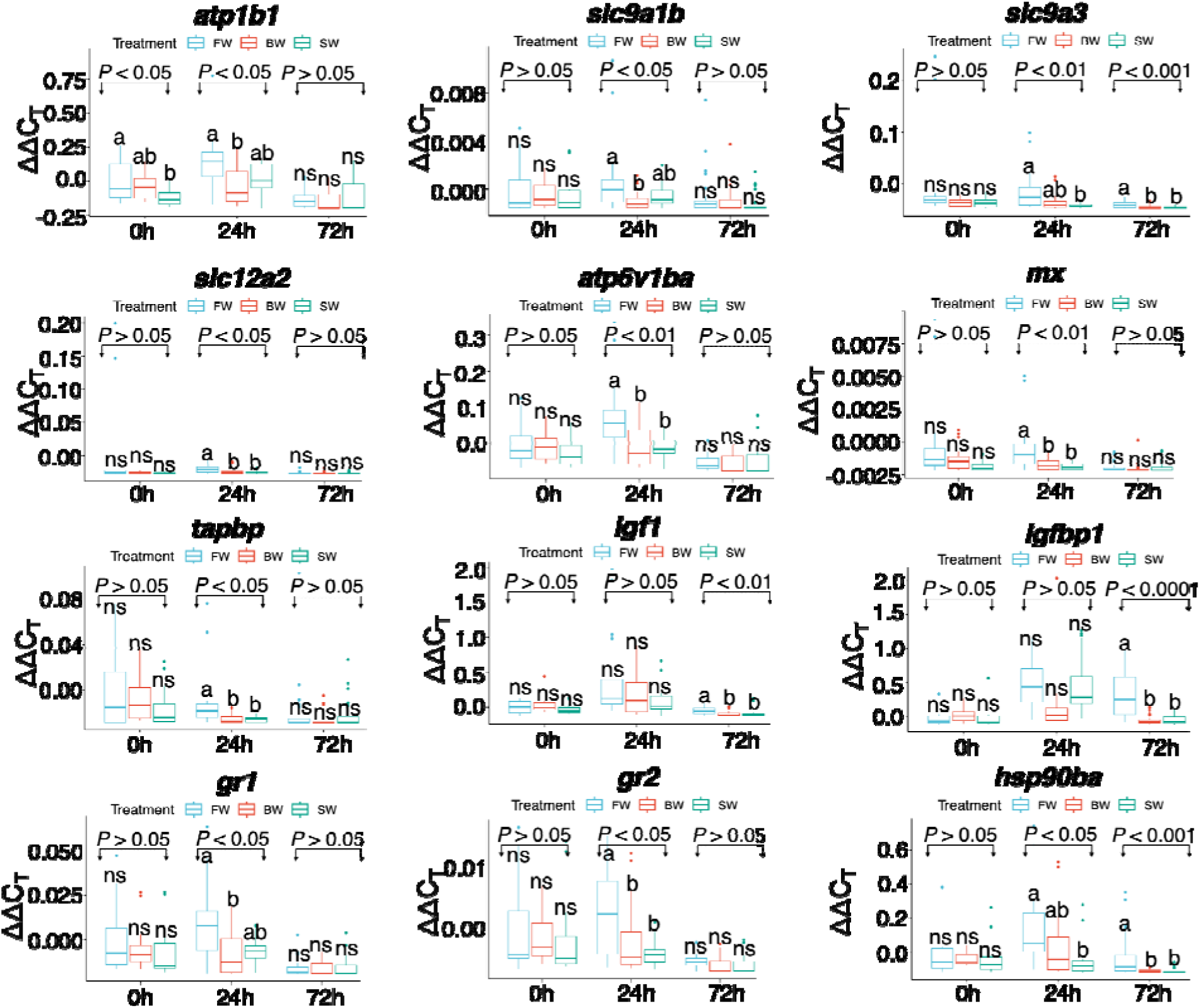
Box plots represent 12 genes that showed significant differences in transcript abundances in liver tissue based on mixed-effects models. The bold line represents the median, the boxes show the 25^th^ and 75^th^ percentiles, the whiskers indicate the 95% confidence interval, and the dots are outliers. Different letters denote significant differences in mean ΔΔC_T_ values among treatments.

**Figure 4:**
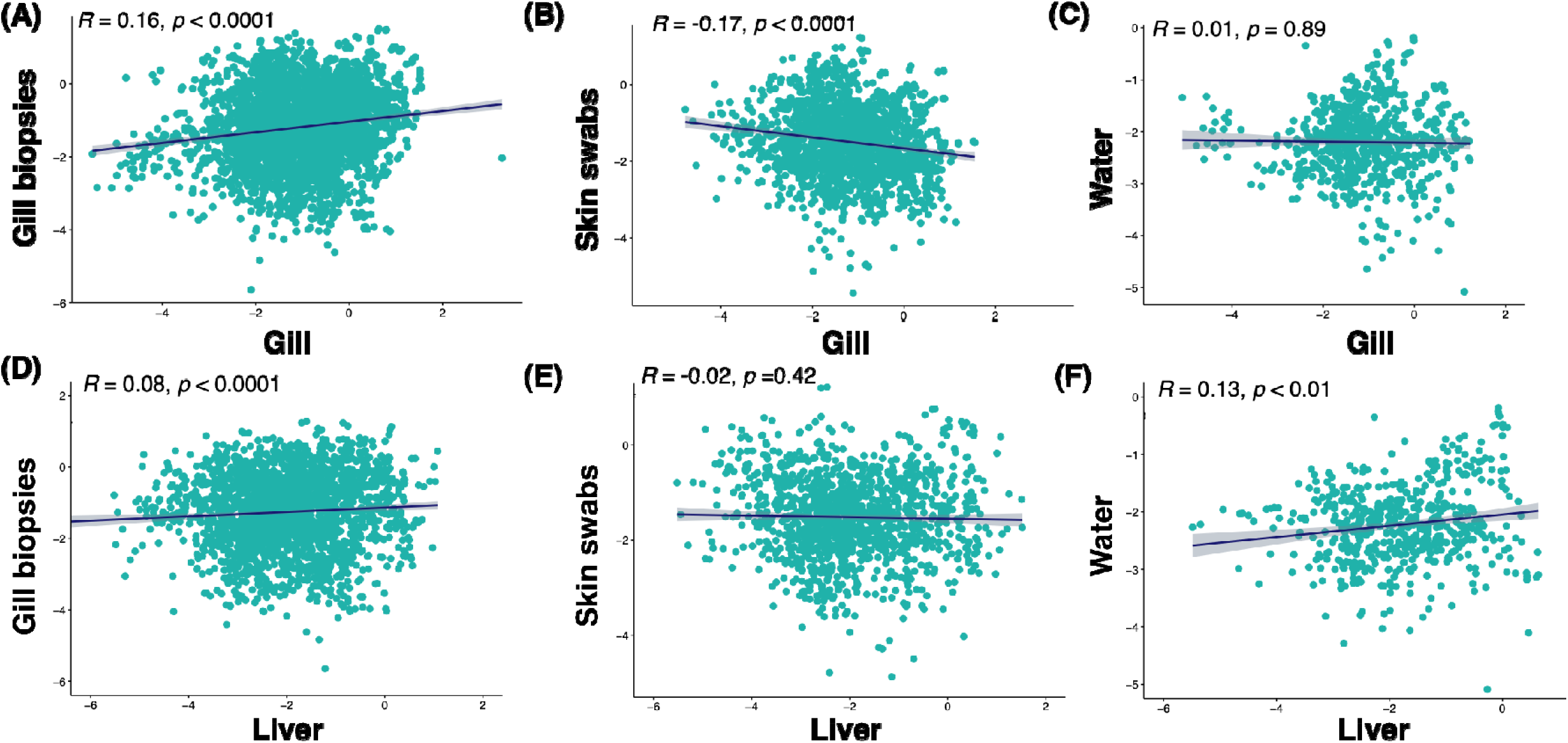
Scatter plots (A-F) represent correlation coefficients (r) between the transcript abundance of the targeted genes between lethal tissue (gill or liver in x-axis) and non-lethal tissue (gill biopsies or skin swabs or water). Regression line indicates strength and direction of relationship.

**Table 2:**
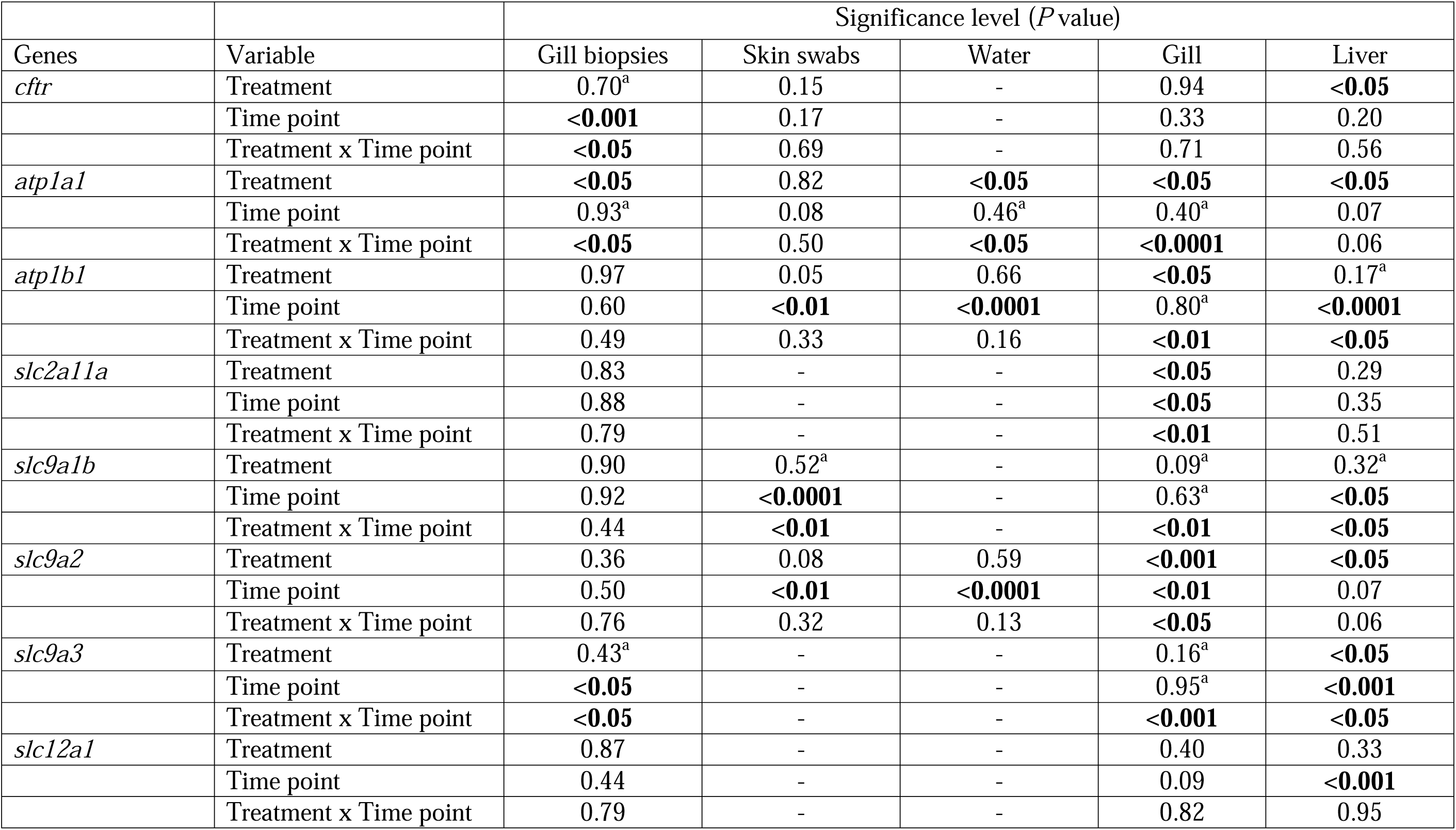
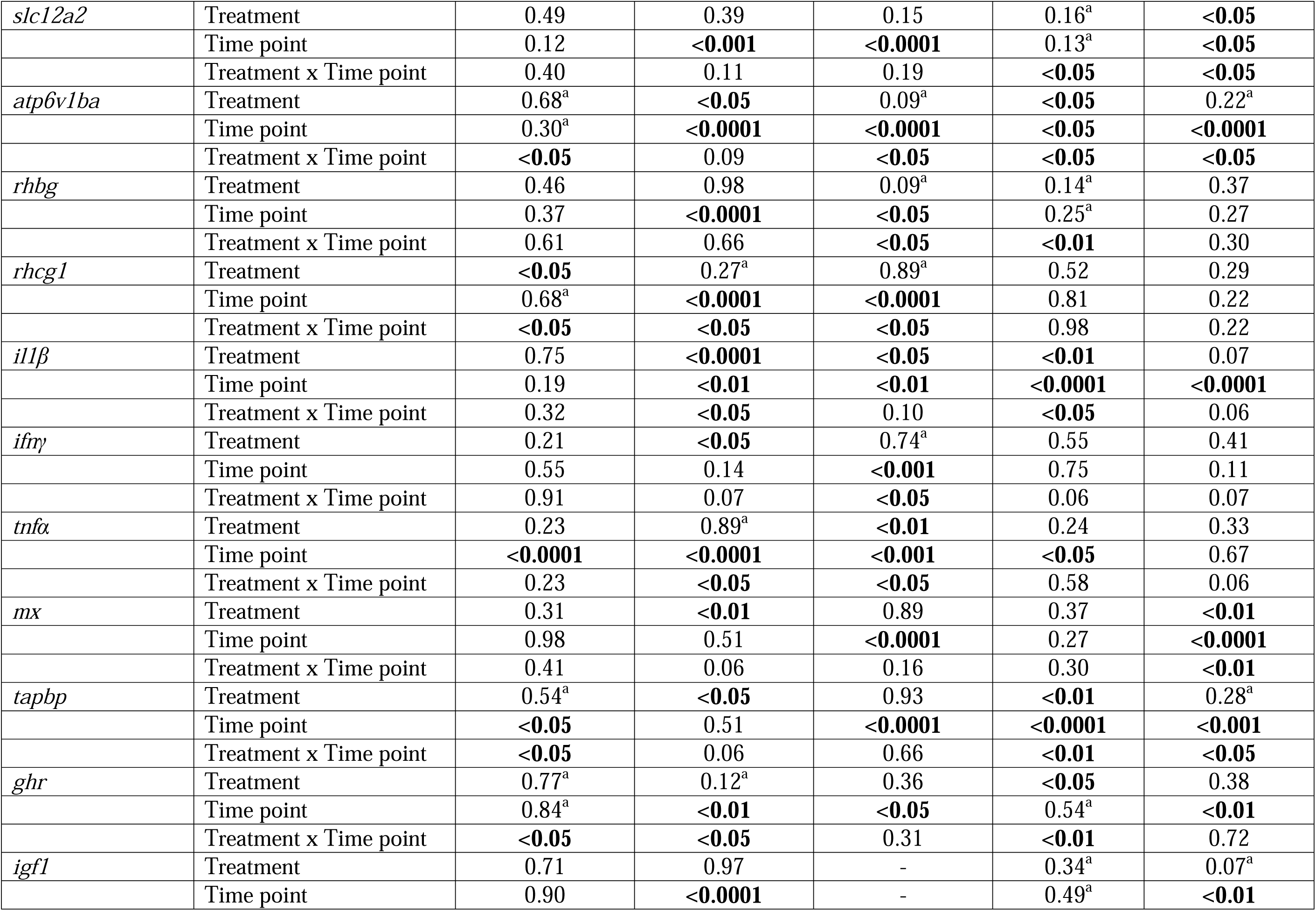
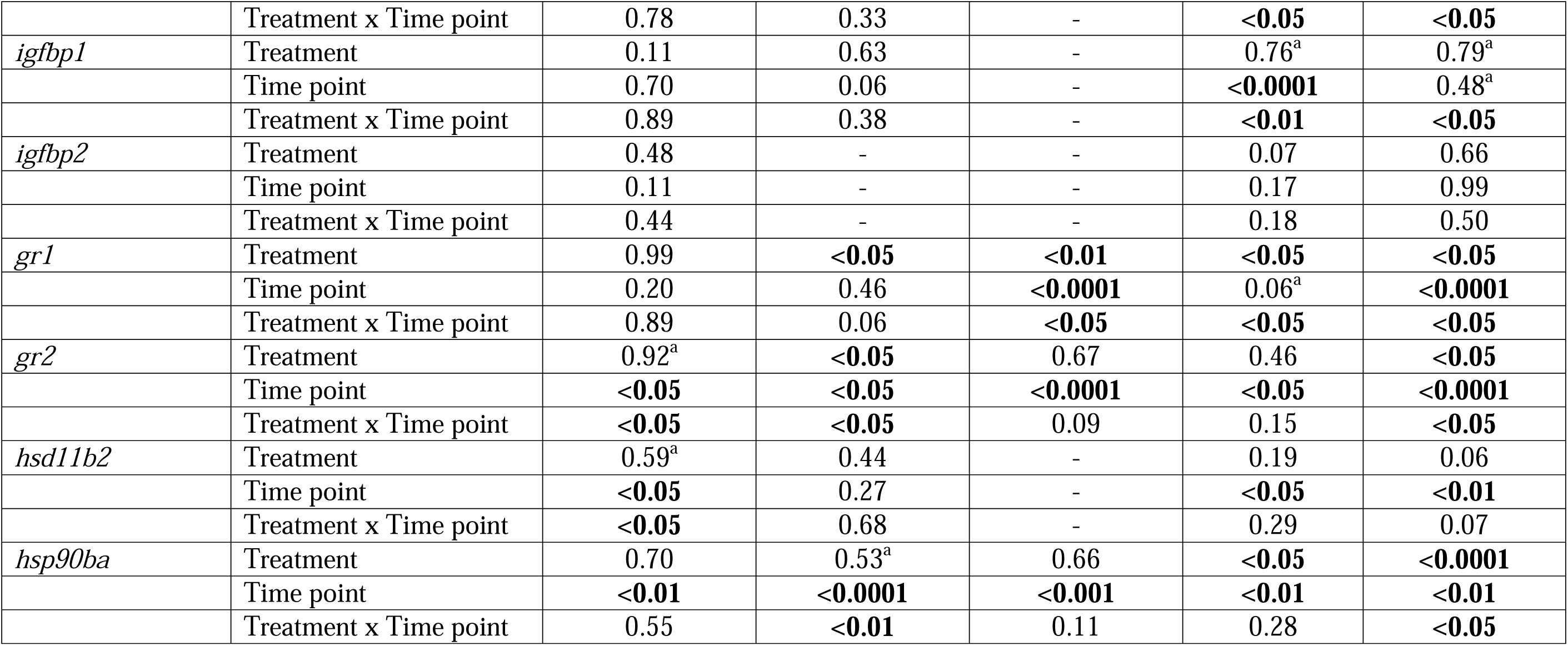
Summary of linear mixed-effects models for transcript abundance differences across tissues (gill biopsies, skin swabs, water, gill, and liver). Treatment and time point were included as fixed effects, and tank ID was included as a random intercept in the model. Significance values ((p-values in **bold**) for the fixed effects and interaction term were obtained using an F-test based on Satterwaite’s approximation. Genes that did not show expression across all treatments and time-points were not included in the model and are presented as “–”. ^a^If the interaction term was significant, insignificant fixed effects were retained in the final model.

## Results

Among non-lethal tissues, 23 assays showed more than 80% amplification success for all treatment-by-time combinations with gill biopsies tissue cDNA, whereas only five assays for skin swabs, and none of the assays for water, showed more than 80% amplification success (Table 1). Among lethal tissues, 25 assays showed more than 80% amplification success for all treatment-by-time combinations with gill tissue, whereas 13 assays for liver tissue showed more than 80% amplification success (Table 1). An 80% amplification success threshold was selected a priori as a criterion to identify assays that amplified consistently across the majority of biological samples for all treatment-by-time combinations. Among the amplified assays in non-lethal tissue samples, nine assays in gill biopsies and seven assays in both skin swabs and water samples showed significant differences in transcript abundance for treatment and/or time point and the treatment x time point interaction and, therefore, were retained in the final LME models (Table 2). In lethal sample tissues, 15 assays in gill (Table 2, Figure 2) and 12 assays in liver tissues (Table 2, Figure 3) showed significant differences in transcript abundance for treatment and/or time point and the treatment x time point interaction.

In general, within non-lethal tissue samples, gill biopsy samples (Supplementary Figure S1) showed no significant differences in transcript abundance among treatments at the 0 h time point. At 24 h and 72 h after exposure, significant differences among three treatments were detected (except for *tapbp* and *ghr* at 24 h). Overall, expression levels were upregulated in SW samples compared to FW samples. For skin swab samples (Supplementary Figure S2), there were also no significant differences in transcript abundance among treatment at the 0 h and 24 h time points (except for *il1*β at 24 h). At 72 h, significant differences in transcript abundance among treatments were also observed. However, in skin swab samples, expression levels were downregulated in SW samples compared to FW samples. For water samples (Supplementary Figure S3), similarly, at the 0 h time point, there were no significant differences in transcript abundance among treatment (except, *atp6v1ba*, *ifn*γ, *tnf*α, and *gr1*). However, significant differences were observed at the 24 h time point (except, *rhcg1*, *tnf*α, and *gr1*). Surprisingly, among seven genes, only *rhcg1* showed significant differences in transcript abundance among treatment at the 72 h time point.

In contrast, within lethal tissue samples, gill samples (Figure 2) showed no significant differences in transcript abundance among treatments at the 0 h (except for *atpa1a*, *slc2a11a*, *atp6v1ba*, and *il1*β) and 24 h (except for *slc9a2*, *slc12a2*, *il1*β, and *igfbp1*) time points. At 72 h after exposure, significant differences in transcript abundance among treatments were observed. Overall, expression levels were upregulated in SW samples compared to control FW samples. For liver tissue (Figure 3), there were no significant differences in transcript abundance among treatments at the 0 h time point (except for *atpa1b*). At 24 h after exposure, significant differences in transcript abundance among treatments were observed, with some exceptions (e.g., *igf1* and *igfbp1*). However, at 72 h after exposure, only four assays (e.g., *slc9a3*, *igf1*, *igfbp1*, and *hsp90ba*) showed significant differences in transcript abundance among treatments. Overall, the expression levels were upregulated in SW samples compared to control FW samples, with a few exceptions.

A PCA of the amplified qPCR genes in both non-lethal and lethal tissues showed distinct clustering across treatments and time points (Supplementary Figure S4). Overall, PC1 explained 48.4%, 32.0%, 27.0%, 75.7%, and 31.2% of the variance in mRNA transcript abundance patterns in gill biopsies, skin swabs, water, gill, and liver tissue samples, respectively, at the 0 h time point. In contrast, PC2 explained 13.9%, 14.4%, 20.2%, 8.8%, and 17.2% of the variance in transcript abundance in gill biopsies, skin swabs, water, gill, and liver tissue samples, respectively, at the 0 h time point. At the 24 h time point, PC1 explained 60.2%, 32.4%, 21.2%, 62.2%, and 44.7% of the variance in gill biopsies, skin swabs, water, gill, and liver tissue samples, respectively, whereas PC2 explained 15.1%, 14.7%, 14.6%, 12.2%, and 7.7% of the variance, respectively. Finally, at the 72 h time point, PC1 explained 64.0%, 32.0%, 30.3%, 69.4%, and 26.0% of the variance in gill biopsies, skin swabs, water, gill, and liver tissue samples, respectively, whereas PC2 explained 9.6%, 15.4%, 14.5%, 6.7%, and 10.5% of the variance, respectively.

The correlation coefficients (R) between the transcript abundance of the targeted genes among lethal and non-lethal tissues were significantly positive, with a few exceptions (Figure 4). For example, the correlation of transcript abundance for non-lethal gill biopsies was significantly positive with both lethal gill (*R* = 0.16, *p* < 0.0001) and liver samples (*R* = 0.08, *p* < 0.0001). Surprisingly, the correlation of transcript abundance of targeted genes for non-lethal skin swab samples was significantly negative with lethal gill samples and showed no significant correlation with liver samples. The water samples were positively correlated (but not significantly so) with lethal gill samples and significantly positively correlated with liver samples.

## Discussion

The present study characterized transcriptomic responses of Chinook salmon smolts to osmoregulatory challenges, a critical process as juveniles migrate downstream to the sea during early life stages. This controlled hatchery-based gene transcriptional profiling study provides insights into Chinook salmon hatchery rearing and release timing, while demonstrating potential future field applications, including monitoring of physiological health of wild juveniles and assessing their responses to climate-related environmental stressors. The main findings of this study can be summarized as: (i) most assays showed higher qPCR amplification success in both non-lethal gill biopsies and lethal gill tissues, but fewer in skin swabs and water (non-lethal) and liver (lethal) samples; (ii) more assays showed significant differences in transcript abundance in traditional lethal tissues compared to non-lethal tissues; (iii) with increasing time of salinity exposure, gene transcript levels were upregulated in stressed SW samples compared to control FW samples for both non-lethal and lethal tissue samples; (iv) targeted genes for both non-lethal and lethal tissues showed distinct clustering across treatment and time points, but no clear association within biological functions; and (v) the correlation coefficients of the targeted genes between non-lethal and lethal tissues were significantly positive, with a few exceptions. Our findings will help foster better husbandry practices by identifying gene transcriptional responses associated with the physiological status of juvenile Chinook salmon, which could be used to evaluate their coping response to hatchery rearing conditions as well as inform decisions related to release timing. Moreover, the application of these bioassays for wild Chinook salmon juveniles could provide valuable insights into their physiological status in relation to environmental stressors, supporting the conservation and management of wild stocks.

In the present study, our qPCR assays showed better amplification success in gill tissues (gill biopsies and gill samples) compared to other non-lethal and lethal tissue samples (McCormick, 1994; Cooke et al., 2005). Fish gills are extremely sensitive to environmental stressors, particularly salinity changes (Maryoung et al., 2015). Moreover, the gill is a multifunctional organ that not only mediates gas exchange but also plays roles in osmotic and ionic regulation, acid-base balance, and ammonia excretion (Evens et al., 2005). Furthermore, gill tissue is not only a target organ for osmoregulatory-relevant genes, but it also plays important roles in other biological functions such as metabolism (Zhao et al., 2021), as well as thermal stress and immune responses (Best et al., 2025; Howarth et al., 2025; Dang et al., 2026). It is not surprising that some qPCR assays did not amplify in non-lethal (skin swabs and water) and lethal (liver), a pattern likely reflecting either low expression of osmoregulatory genes (e.g., liver samples) or low transcript recovery (e.g. water samples). Finally, it is likely that our qPCR assays did not perform well in non-gill tissue samples because these osmoregulatory-relevant genes were specifically targeted for gill tissues.

One of the striking findings of this current study was that more assays showed significant differences in transcript abundance in traditional lethal tissues (both gill and liver) compared to non-lethal tissues (gill biopsies, skin swabs, and water samples). It is evident that gene transcription induced by salinity change is highly tissue-specific (Seear et al., 2010; Maryoung et al., 2015). It has previously been suggested that different organs and cell types within an organism may detect and respond differently to changes in osmolality (Fiol & Kültz, 2007). Accordingly, internal organs such as the liver may exhibit different transcriptional responses to salinity changes than non-lethal sampled tissues such as skin swabs, while the transcript abundance detectable in the surrounding water may also vary with salinity. Surprisingly, the targeted genes that showed significant differences in transcript abundances were not entirely consistent between non-lethal gill biopsies and lethal gill samples. For example, in non-lethal gill biopsy tissue, *cftr*, *atpa1a*, *slc9a3*, *atp6v1ba*, *rhcg1*, *tapbp*, *ghr*, *gr2*, *and hsd11b2* showed significant differences in transcript abundance, whereas, in lethal gill tissue, in addition to similar candidate genes (*atpa1a*, *slc9a3*, *atp6v1ba*, *tapbp*, and *ghr*), *atpa1b*, *slc2a11a*, *slc9a1b*, *slc9a2*, *slc12a2*, *rhbg*, *il1*β, *igf1*, igfbp1, and *gr1* also showed significant differences in transcript abundance. This difference is likely due to variation in in the tissue composition of the two sample types. For example, non-lethal gill biopsies consisted only of the distal tips of the gill filaments, whereas lethal gill samples included gill filaments, gill rakers, and the gill arch. As these gill structures differ in their cellular and physiological functions, the extracted RNA may result in differences in the transcript abundances detected in these two gill sampling methods.

Another key finding of this study was that, even with 72 h acute salinity challenge, gene transcript levels were upregulated in stressed SW samples compared to control FW samples in both non-lethal and lethal tissue samples. It is well documented that juvenile Chinook salmon migration from fresh water to sea water during early life stages is a critical process involving substantial changes in behaviour, skin pigmentation, and body morphology (McCormick et al., 1998, 2013). The physiological changes during migration are equally important, including shifts in hemoglobin isoforms, increased metabolic rate and enhanced seawater tolerance, which are essential for immediate survival (Robertson & McCormick, 2012). Undeniably, juvenile Chinook salmon that are unprepared for increased salinity, remain in fresh water too long, or are otherwise poorly suited to seawater conditions exhibit greatly reduced survival and slower growth due to ionic and osmotic imbalance caused by the higher ion concentrations in seawater relative to freshwater. Even with only a short window of salinity exposure (72 h), significant upregulation of targeted gene transcripts was detected in SW-exposed fish compared with FW control fish across both non-lethal and lethal tissue samples (with a few exceptions), indicating a rapid transcriptional response to increased salinity. We did not observe any smolt mortalities during the exposure time. Moreover, despite the short duration of our salinity exposure, our transcriptomic tools detected transcriptional responses to salinity under controlled hatchery conditions, providing their potential for field applications to assess the physiological status of juvenile Chinook salmon and their responses to changing environments using non-lethal sampling approaches.

Lastly, for both non-lethal and lethal samples, the lack of clustering of targeted genes according to their biological functions suggests that the transcriptional response of juvenile Chinook salmon to salinity is not restricted to individual physiological and molecular pathways, but involves responses across multiple biological processes. This is consistent with the complex physiological adjustments that juvenile Chinook salmon undergo during the transition from fresh water to sea water, during which different biological processes associated with osmoregulation, growth and metabolism, stress response, immune function may respond simultaneously and interact with one another. Furthermore, many genes have roles in multiple biological functions, which may explain why genes assigned to different functional categories exhibited similar transcriptional patterns across treatments and time points (Islam et al., 2026). Moreover, the predominantly positive correlations among targeted genes between non-lethal and lethal tissues further suggest coordinated transcriptional responses during salinity challenges (Finerty et al., 2026). In contrast, the significant negative correlation between non-lethal skin swab and lethal gill samples may reflect differences in the cellular functions represented by these sample types, indicating they may capture different physiological responses to changes in salinity.

Chinook salmon play a vital ecological, economic, and societal role in Western Canada (BC). However, under anthropogenic influences such as climate change, Chinook salmon stocks are under threat. To mitigate these threats, effective conservation and management approaches are needed for Chinook salmon populations in BC. Currently, logistical difficulties in monitoring the physiological health status of Chinook salmon populations are compounded by the limitations of conventional captured based methods. Transcriptional profiling using non-lethal tissue samples, such as gill biopsies and skin swabs can provide information on the physiological status of juvenile Chinook salmon, while avoiding the need for lethal sampling. Furthermore, the detection of transcript abundance in water samples (eRNA), if further validated under field conditions, could provide an additional layer for assessing physiological responses while avoiding direct fish capture or even contact. The chip used in this study could be used as a tool to measure transcriptional profiling responses of juvenile Chinook salmon of fisheries and aquaculture importance in BC to known osmoregulatory challenges. This work demonstrates that an osmoregulatory chip-based transcriptional profiling approach could be widely adopted for monitoring health in hatchery/aquaculture and wild Chinook salmon populations.

## Acknowledgements

The authors would like to thank Dr Ann Heath and staffs of Yellow Island Aquaculture Ltd. for technical assistance. Yellow Island Aquaculture Ltd provided logistical and in-kind support for this project.

## Author Contributions

SSI contributed to study design, completed the lab work, data analyses and manuscript preparation. KMJ, JWH, and DDH contributed to study design, data analyses and manuscript preparation and revision. KMJ and DDH contributed to secure funding.

## Data Availability

The qPCR metadata for gill biopsies, skin swabs, water, gill, and liver tissues across treatments and time points for multivariate analyses (see Supplementary file) have been submitted to Figshare: 10.6084/m9.figshare.33158483

## Ethics Statement

All animals were treated following the guidelines provided by the Canadian Council on Animal Care during holding and experimentation, and approval was granted by the University of Windsor Animal Utilization Project Proposal (AUPP#23-06). Animals were euthanized with an overdose of clove oil (Aromatics Canada Inc; 0.4 mL/L clove oil in aerated hatchery water).

