## Supplementary File for "Gene transcriptional profiling to assess Chinook salmon osmoregulatory resilience to salinity challenges"

Supplementary Table S1: Fork length (cm) and weight (g) of individual fish used for molecular analyses in this study.

| **Treatment** | **Time point** | **Tank ID** | **Fish ID** | **Fork length (cm)** | **Weight (g)** |
| --- | --- | --- | --- | --- | --- |
| FW | 0 | 1 | 1 | 16 | 48.1 |
| FW | 0 | 1 | 2 | 15.7 | 50.8 |
| FW | 0 | 2 | 1 | 14.2 | 35.4 |
| FW | 0 | 2 | 2 | 14.5 | 38.1 |
| FW | 0 | 3 | 1 | 16 | 47.2 |
| FW | 0 | 3 | 2 | 14.6 | 36.3 |
| FW | 0 | 4 | 1 | 15.5 | 45.4 |
| FW | 0 | 4 | 2 | 17 | 59.9 |
| BW | 0 | 5 | 1 | 14.5 | 36.3 |
| BW | 0 | 5 | 2 | 16.3 | 50.8 |
| BW | 0 | 6 | 1 | 16 | 48.1 |
| BW | 0 | 6 | 2 | 16.5 | 52.6 |
| BW | 0 | 7 | 1 | 15 | 39.9 |
| BW | 0 | 7 | 2 | 15.5 | 49.0 |
| BW | 0 | 8 | 1 | 14 | 30.8 |
| BW | 0 | 8 | 2 | 16 | 48.1 |
| SW | 0 | 9 | 1 | 15 | 40.8 |
| SW | 0 | 9 | 2 | 14.5 | 36.3 |
| SW | 0 | 10 | 1 | 14 | 31.8 |
| SW | 0 | 10 | 2 | 14 | 32.7 |
| SW | 0 | 11 | 1 | 16.5 | 53.5 |
| SW | 0 | 11 | 2 | 15 | 44.5 |
| SW | 0 | 12 | 1 | 15 | 40.8 |
| SW | 0 | 12 | 2 | 14.5 | 35.4 |
| FW | 24 | 1 | 1 | 15 | 38.1 |
| FW | 24 | 1 | 2 | 15.7 | 40.8 |
| FW | 24 | 2 | 1 | 15.8 | 47.2 |
| FW | 24 | 2 | 2 | 15.5 | 47.2 |
| FW | 24 | 3 | 1 | 16.5 | 49.0 |
| FW | 24 | 3 | 2 | 14.5 | 36.3 |
| FW | 24 | 4 | 1 | 16.5 | 56.2 |
| FW | 24 | 4 | 2 | 17 | 62.6 |
| BW | 24 | 5 | 1 | 17 | 59.0 |
| BW | 24 | 5 | 2 | 16 | 49.0 |
| BW | 24 | 6 | 1 | 16 | 51.7 |
| BW | 24 | 6 | 2 | 15.5 | 47.2 |
| BW | 24 | 7 | 1 | 14.9 | 41.7 |
| BW | 24 | 7 | 2 | 14.8 | 40.8 |
| BW | 24 | 8 | 1 | 16.8 | 52.6 |
| BW | 24 | 8 | 2 | 14.5 | 38.1 |
| SW | 24 | 9 | 1 | 14.8 | 38.1 |
| SW | 24 | 9 | 2 | 14.8 | 33.6 |
| SW | 24 | 10 | 1 | 14.5 | 36.3 |
| SW | 24 | 10 | 2 | 14.5 | 34.5 |
| SW | 24 | 11 | 1 | 15.5 | 47.2 |
| SW | 24 | 11 | 2 | 16.3 | 53.5 |
| SW | 24 | 12 | 1 | 16.5 | 56.2 |
| SW | 24 | 12 | 2 | 14.5 | 36.3 |
| FW | 72 | 1 | 1 | 16.8 | 49.9 |
| FW | 72 | 1 | 2 | 16.5 | 55.3 |
| FW | 72 | 2 | 1 | 15.5 | 43.5 |
| FW | 72 | 2 | 2 | 16.5 | 53.5 |
| FW | 72 | 3 | 1 | 15.5 | 42.6 |
| FW | 72 | 3 | 2 | 16.4 | 51.7 |
| FW | 72 | 4 | 1 | 15.5 | 43.5 |
| FW | 72 | 4 | 2 | 15.3 | 43.5 |
| BW | 72 | 5 | 1 | 15 | 38.1 |
| BW | 72 | 5 | 2 | 15.7 | 43.5 |
| BW | 72 | 6 | 1 | 16 | 48.1 |
| BW | 72 | 6 | 2 | 16.5 | 52.6 |
| BW | 72 | 7 | 1 | 14.9 | 40.8 |
| BW | 72 | 7 | 2 | 16 | 49.0 |
| BW | 72 | 8 | 1 | 15.5 | 48.1 |
| BW | 72 | 8 | 2 | 16.3 | 50.8 |
| SW | 72 | 9 | 1 | 15.5 | 39.0 |
| SW | 72 | 9 | 2 | 15.6 | 49.0 |
| SW | 72 | 10 | 1 | 15.4 | 38.1 |
| SW | 72 | 10 | 2 | 16.2 | 45.8 |
| SW | 72 | 11 | 1 | 16 | 55.3 |
| SW | 72 | 11 | 2 | 15.3 | 41.3 |
| SW | 72 | 12 | 1 | 17.5 | 63.5 |
| SW | 72 | 12 | 2 | 14.3 | 36.3 |

Supplementary Table S2: Osmoregulatory stress-response transcriptional profiling nanofluidic chip candidate genes used for Taqman^TM^ qPCR assays of gene transcription on the OpenArray® platform. The candidate genes are arranged by putative function, and gene name (symbol), forward and reverse primer and probe sequences, amplicon size, accession numbers and species used in the final alignments for primer design are listed.

| Gene Function | Gene Name | Forward/Reverse Primers & Probe sequence | Amplicon size (bp) | Species used in the final alignments for primer design | Accession Number |
| --- | --- | --- | --- | --- | --- |
| Osmoregulation | Cystic fibrosis transmembrane conductance regulator (Cl- channel)  (*cftr*) | F: CAGGCAGAAAAGAGCAGGGA  R: TCTCCATCACCTCCTCCCAG  P: AGTCGTCGCCTGGC | 107 | *S. salar*  *S. trutta*  *O. keta*  *O. mykiss*  *O. tshawytscha*  *O. nerka*  *S. namaycush*  *S. alpinus*  *C. clupeaformis* | NM_001123533  XM_029697084  XM_035798038  XM_036979278  XM_042323330  XM_029634857  XM_039000654  XM_023970781  XM_041836795 |
|  | Sodium-potassium ATPase alpha subunit (*atp1a1*) | F: TGTGTCCGTCTTCTTCGGTG  R: CAGTAGCCAGGAGACCCTCTG  P: GCTATGGATGGCTGCC | 126 | *O. mykiss*  *S. salar*  *S. trutta*  *O. tshawytscha*  *O. kisutch*  *O. masou*  *S. namaycush* | NM_001124461  XM_045722983  XM_029748747  XM_042316867  XM_031818113  AB573640  XM_038981298 |
|  | Sodium-potassium ATPase subunit beta-1 (*atp1b1*) | F: GGATGCTTGGCTGGGATTT  R: CTCTGTCCTGGTAGGTGGGTTT  P: CAGGCCCTGCTGCT | 83 | *S. salar*  *S. trutta*  *O. mykiss*  *O. tshawytscha*  *O. kisutch*  *S. alpinus*  *S. namaycush*  *C. clupeaformis* | NM_001252362  XM_029708332  XM_036979818  XM_042322829  XM_031822295  XM_024142989  XM_039003039  XM_041840719 |
|  | Solute carrier family 2-member 11a  (*slc2a11a*) | F: GTTCGCTGACCGTAATGTGC  R: ATGGAGTCGTTGCCACAGAG  P: CAGCAGCGCCATGC | 84 | *S. salar*  *O. keta*  *O. mykiss*  *O. tshawytscha*  *O. kisutch*  *O. gorbuscha*  *S. namaycush*  *S. alpinus*  *C. clupeaformis* | NM_001141793  XM_035778241  XM_036950146  XM_024374110  XM_020486753  XM_046311987  XM_038970559  XM_071380116  XM_041889167 |
|  | Solute carrier family 9 member A1B  (*slc9a1b*) | F: CTTCATCTTCCTGGGCGTGT  R: GACGAAGGTCCAGTTCCAGG  P: CACGGTGGCAGGGC | 61 | *O. mykiss*  *S. salar*  *O. keta*  *O. tshawytscha*  *O. gorbuscha*  *O. kisutch*  *S. namaycush* | NM_001124695  XM_014142630  XM_035757656  XM_024395440  XM_046361737  XM_020500134  XM_038972353 |
|  | Solute carrier family 9 member 2 (*slc9a2*) | F: CATCTCAGCTGTGGACCCAG  R: TCGTTGAACAGGCACTCTCC  P: CCGTGCTCAGCGTGT | 102 | *O. mykiss*  *S. salar*  *S. trutta*  *O. tshawytscha*  *O. nerka*  *O. kisutch*  *S. alpinus*  *S. namaycush*  *C. clupeaformis* | NM_001130994  XM_014174391  XM_029712099  XM_024410130  XM_029646675  XM_020499911  XM_024010456  XM_039006593  XM_041843915 |
|  | Solute carrier family 9 member A3, tandem duplicate 2  (*slc9a3*) | F: ATCTTCGTCCTGGGCTACCTC  R: ACAGCAGACCCCACAGAATG  P: AGCCGAGATGCTCT | 90 | *O. mykiss*  *S. salar*  *S. trutta*  *O. keta*  *O. tshawytscha*  *O. gorbuscha*  *O. nerka*  *O. kisutch*  *S. alpinus* | NM_001160482  XM_014141359  XM_029712099  XM_035793565  XM_024396240  XM_046362575  XM_029663816  XM_020500716  XM_023975187 |
|  | Solute carrier family 12 member 1  (*slc12a1*) | F: CCTGGTTGGACATCCTCACC  R: GAAGGTGAGGACGCTCTTGTG  P: ACCTCCCTCCCACCAT | 77 | *O. tshawytscha*  *S. salar*  *S. trutta*  *O. mykiss*  *O. kisutch*  *S. namaycush* | XM_042316068  NM_001141520  XM_029720918  XM_036981072  XM_031820808  XM_038987605 |
|  | Solute carrier family 12 member 2 (Na-K-Cl cotransporter isoform 1)  (*slc12a2*) | F: CGTTTTGCATCGCTCTAGCC  R: AGCATAGGAGGCCAGGAAGA  P: TCGCTCCCATCATCT | 89 | *S. salar*  *S. trutta*  *O. keta*  *O. mykiss*  *O. tshawytscha*  *O. nerka*  *O. gorbuscha*  *S. alpinus*  *S. namaycush* | NM_001123683  XM_029710291  XM_031802486  XM_036976792  XM_042302639  XM_029647200  XM_046290487  XM_023984575  XM_038970029 |
|  | ATPase, H+ transporting, lysosomal, V1 subunit B, member a  (*atp6v1ba*) | F: AGAAGACGGCCTGTGAGTTCA  R: GTCCCAGCATGTCCTCAGACA  P: CGGTGACATCCTGCGC | 66 | *S. salar*  *O. mykiss*  *O. tshawytscha*  *S. namaycush* | NM_001141746  XM_021609923  XM_024424712  XM_038960498 |
|  | Rh family, B glycoprotein  (*rhbg*) | F: CACAGCCACCGTTCTGATCTC  R: TGATGAGGAGCTGGACAGGAC  P: CGTTCGGGGCAGTGC | 68 | *S. salar*  *S. trutta*  *O. keta*  *O. tshawytscha*  *O. nerka*  *O. gorbuscha*  *O. kisutch*  *S. alpinus* | NM_001141124  XM_029756072  XM_035774348  XM_024377707  XM_029677301  XM_046297897  XM_020455545  XM_023975561 |
|  | Rh type C glycoprotein  (*rhcg1*) | F: TTCCTGTGGATGTTCTGGCC  R: AGGTGTTGATGACTGCCCTG  P: CATGGAGACGGTCAGC | 82 | *O. mykiss*  *S. salar*  *S. trutta*  *O. keta*  *O. tshawytscha*  *O. gorbuscha*  *O. nerka*  *O. kisutch*  *S. alpinus* | NM_001124523  XM_014175744  XM_029721355  XM_035787123  XM_024440305  XM_046345896  XM_029639391  XM_031812239  XM_024002622 |
| Immune  Function | Interleukin 1β (*il-1β*) | F: ACCTGTCCTGCTCCAAATCG  R: TGCTGGCTGATGGACTTCAG  P: CCCACCCTGCACCT | 88 | *S. salar*  *S. trutta*  *O. keta*  *O. mykiss*  *O. nerka*  *O. tshawytscha*  *O. kisutch*  *S. alpinus*  *S. namaycush*  *C. clupeaformis* | NM_001123582  XM_029716234  XM_035786315  NM_001124347  XM_029638479  XM_024418276  XM_020475860  XM_024002415  XM_039003225  XM_041901577 |
|  | Interferon γ (*ifnγ*) | F: GGAAGGCTCTGTCCGAGTTC  R: CTCTTCCTGCGGTTGTCCTT  P: CCTCCAAACTGGCC | 91 | *S. salar*  *S. trutta*  *O. mykiss*  *O. tshawytscha*  *O. namaycush*  *O. kisutch*  *C. clupeaformis* | NM_001171804  XM_029743588  FM864345  XM_042314919  XM_038995696  XM_031819098  XM_041873105 |
|  | Tumor necrosis factor α (*tnf-α*) | F: AGTACCAGACGCTGCTCAACTC  R: TTTTCCTTCACTCGCAGCCT  P: TCTGTACGCACCGTGTGT | 77 | *S. salar*  *S. trutta*  *O. keta*  *O. mykiss*  *O. tshawytscha*  *O. gorbuscha*  *O. kisutch*  *S. alpinus*  *S. namaycush* | NM_001123617  XM_029734485  XM_035767308  HE798146  XM_024407165  XM_046360750  XM_020468340  XM_024139369  XM_038972209 |
|  | Mx protein (*mx*) | F: GGAGGAGATTGAGGACCCCT  R: ATCACTGATACCCACCCCCA  P: TGAAGCCCAGGATGAA | 88 | *S. salar*  *S. trutta*  *O. keta*  *O. mykiss*  *O. nerka*  *O. tshawytscha*  *O. gorbuscha*  *O. kisutch*  *S. alpinus*  *S. namaycush* | NM_001123690  XM_029774982  XM_035777514  XM_021569609  XM_029683585  XM_024415945  XM_046313204  XM_020491468  XM_023993827  XM_039009004 |
|  | Tapasin  (*tapbp*) | F: ACGGCAAGACTGACCGATTT  R: CACTTCAGCTTCCTGCAGGAT  P: AGCGGGGCTAGACTT | 111 | *S. salar*  *S. trutta*  *O. keta*  *O. mykiss*  *O. nerka*  *O. tshawytscha*  *O. gorbuscha*  *O. kisutch*  *S. alpinus*  *S. namaycush*  *C. clupeaformis* | NM_001123605  XM_029731732  XM_035795285  NM_001124553  DQ092324  XM_024441639  XM_046326692  XM_020505550  XM_023976454  XM_038966528  XM_041903149 |
| Growth and Metabolism | Growth hormone receptor  *(ghr)* | F: TAACCGGGAGCCACTTTGAC  R: ATTGACCTCACGGTACTGCACC  P: TGAGCTGGGAGCCG | 110 | *S. salar*  *S. trutta*  *O. keta*  *O. mykiss*  *O. nerka*  *O. tshawytscha*  *O. gorbuscha*  *O. kisutch*  *O. masou*  *S. alpinus*  *S. namaycush*  *C. clupeaformis* | NM_001123594  XM_029769265  XM_035772405  NM_001124731  XM_029620369  XM_024397386  XM_046317034  AF403540  AB071216  XM_023987611  XM_038964896  XM_041850797 |
|  | Insulin-like growth factor 1  (*igf1*) | F: TTCAAGAGTGCGATGTGCTGT  R: CGCCGAAGTCAGGGTTAGG  P: ACACCCTCTCACTGCT | 79 | *S. salar*  *S. trutta*  *O. keta*  *O. mykiss*  *O. nerka*  *O. tshawytscha*  *O. gorbuscha*  *O. kisutch*  *S. alpinus*  *S. namaycush*  *C. clupeaformis* | NM_001123623  XM_029742642  XM_035748910  XM_021577176  XM_029665859  XM_024372796  XM_046297130  XM_031797471  XM_024014490  XM_038977529  XM_045214404 |
|  | Insulin-like growth factor binding protein 1  (*igfbp1)* | F: AGAGCATCAAGGCCAAGGTC  R: CATGCAGCTCAATGTGGCAG  P: ACTGGTGGAACAGGGT | 75 | *S. salar*  *S. trutta*  *O. mykiss*  *O. keta*  *O. tshawytscha*  *O. nerka*  *O. gorbuscha*  *O. kisutch*  *S. alpinus* | NM_001279137  XM_029750591  NM_001124561  XM_035736623  XM_024420224  XM_029687052  XM_046351830  XM_020492902  XM_023997977 |
|  | Insulin-like growth factor binding protein 2  (*igfbp2)* | F: CCTTCACGGGTGTCCAGATC  R: CTCCTCCACATACTGGCTGC  P: CCACGAAGGTGCGGG | 74 | *S. salar*  *S. trutta*  *O. keta*  *O. mykiss*  *O. nerka*  *O. gorbuscha*  *O. kisutch*  *S. alpinus*  *S. namaycush* | NM_001279160  XM_029703049  XM_035774215  XM_021579985  XM_029635193  XM_046368859  XM_020462099  XM_023980724  XM_039014863 |
| Stress  response | Glucocorticoid receptor 1  (*gr1)* | F: GACTCTGCTGCAGTGTTCCTG  R: CACTGGTCTGCCATGTAGGG  P: CCTTCGGCCTGGGC | 150 | *O. mykiss,*  *S. salar*  *O. tshawytscha*  *O. kisutch*  *S. alpinus* | NM_001124482  XM_014198677  XM_042303411  XM_031797047  XM_024137659 |
|  | Glucocorti-coid receptor 2  (*gr2*) | F: ATGGCAGACCAGTGTGAACAGAT  R: GAGCAGCAGCAGAACCTTCAT  P: AGACTGCAGGTGTCTCA | 103 | *O. mykiss*  *S. salar*  *S. trutta*  *O. tshawytscha*  *O. nerka*  *O. kisutch*  *S. alpinus* | AY495372  XM_014198677  XM_029771923  XM_024409462  XM_029655477  XM_031789624  XM_024137659 |
|  | 11β-hydroxysteroid dehydrogenase 2  (*hsd11b2*) | F: CTGTCTAGCAGCGTACGGAGC  R: ATGGTGGACACTTTGACCCC  P: CCTGTTCATCAACACACT | 99 | *O. mykiss*  *S. salar*  *S. trutta*  *O. nerka*  *O. kisutch*  *O. tshawyscha*  *C. clupeaformis* | NM_001124218  XM_014186204  XM_029768326  XM_029651578  XM_031812548  XM_024410377  XM_041859559 |
|  | Heat shock protein 90 (constitutive)  (*hsp90ba*) | F: GAGGTGGAGGAGGACGAGTACA  R: GCTGTGAAGTGGATGTGGGA  P: TCTCCAGGGACACAGAC | 86 | *S. salar*  *S. trutta*  *O. keta*  *O. mykiss*  *O. nerka*  *O. tshawytscha*  *O. kisutch*  *S. alpinus*  *C. clupeaformis* | XM_014208111  XM_029761271  XM_035784415  XM_021577266  XM_029665926  XM_024416091  XM_020491013  XM_023966868  XM_041853504 |
| Reference Genes | Glucose-6-phosphate 1-dehydrogenase  (*g6pd*) | F: GAGTGCCTTCTCCAACCTCCA  R: CAGCCCTTGGTACTCATGCAG  P: CTACCTCGCCCTGCC | 147 | *S. salar*  *S. trutta*  *O. keta*  *O. mykiss*  *O. tshawytscha*  *O. gorbuscha*  *O. kisutch*  *S. namaycush* | NM_001141724  XM_029723525  XM_035738273  XM_021566625  XM_024384803  XM_046365258  XM_020459406  XM_039011009 |
|  | Elongation factor 1 alpha  (*ef1a*) | F: GAAGCTTGAGGACAACCCCA  R: GAAGCTCTCCACACACATGGG  P: CCGCCATCATCGTCAT | 92 | *S. salar*  *S. trutta*  *O. keta*  *O. mykiss*  *O. nerka*  *O. tshawytscha*  *O. kisutch*  *S. alpinus*  *S. namaycush*  *C. clupeaformis* | NM_001141909  XM_029735646  XM_035795162  XM_021561980  XM_029658054  XM_024396037  XM_031793751  XM_024146326  XM_038971575  XM_041903035 |
|  | Ribosomal protein L13a  (*rpl13a*) | F: CACTGGAGAGGCTGAAGGTGT  R: GTGGGCTTCAGACGGACAAT  P: CATGGTCGTACCTGCT | 103 | *O. masou*  *S. salar*  *S. trutta*  *O. keta*  *O. mykiss*  *O. nerka*  *O. tshawytscha*  *O. kisutch*  *S. alpinus*  *S. namaycush*  *C. clupeaformis* | EU325865  XM_014128281  XM_029730684  XM_035742484  XM_021574753  XM_029688610  XM_024437735  XM_020488314  XM_023994886  XM_039009297  XM_041855521 |

Supplementary Table S3: qPCR efficiencies (geometric mean across all time points) for the 28 selected genes in non-lethal (gill biopsy, skin swab, and water) and lethal tissue samples (gill and liver). A “–” denotes that the assay did not amplify in any samples of the corresponding tissue type.

| **Gene symbol** | **qPCR efficiencies (%)** | | | | |
| --- | --- | --- | --- | --- | --- |
|  | **Gill biopsy** | **Skin swab** | **Water** | **Gill** | **Liver** |
| *cftr* | 101.8 | 95.9 | 95.2 | 102.5 | 102.8 |
| *atp1a1* | 114.8 | 105.8 | 112.3 | 113.8 | 116.5 |
| *atp1b1* | 100.8 | 96.0 | 95.5 | 101.3 | 101.7 |
| *slc2a11a* | 96.4 | 95.1 | 95.0 | 97.7 | 98.0 |
| *slc9a1b* | 99.5 | 92.8 | 95.3 | 100.9 | 100.9 |
| *slc9a2* | 98.4 | 93.8 | 96.0 | 100.1 | 100.5 |
| *slc9a3* | 101.3 | 103.5 | 104.3 | 101.7 | 102.0 |
| *slc12a1* | 103.5 | 85.5 | 85.5 | 104.3 | 105.9 |
| *slc12a2* | 100.2 | 96.5 | 94.5 | 101.0 | 101.3 |
| *atp6v1ba* | 98.0 | 96.0 | 93.5 | 99.3 | 99.3 |
| *rhbg* | 96.8 | 95.0 | 92.5 | 97.1 | 98.0 |
| *rhcg1* | 108.7 | 98.5 | 98.8 | 110.3 | 111.5 |
| *il1β* | 94.1 | 92.8 | 93.5 | 95.5 | 95.1 |
| *ifnγ* | 97.1 | 94.5 | 94.8 | 98.1 | 98.5 |
| *tnfα* | 94.3 | 83.7 | 89.5 | 101.3 | 99.0 |
| *mx* | 105.9 | 93.4 | 94.0 | 108.1 | 106.6 |
| *tapbp* | 94.0 | 95.3 | 97.0 | 96.4 | 95.0 |
| *ghr* | 100.5 | 95.5 | 95.0 | 100.5 | 101.1 |
| *igf1* | 100.1 | 92.8 | 91.5 | 99.0 | 100.1 |
| *igfbp1* | 102.0 | 90.3 | 86.0 | 100.6 | 102.4 |
| *igfbp2* | 87.5 | - | - | 94.9 | 92.0 |
| *gr1* | 98.6 | 96.3 | 93.8 | 99.6 | 100.1 |
| *gr2* | 104.9 | 100.5 | 99.5 | 106.0 | 108.2 |
| *hsd11b2* | 99.6 | 90.5 | - | 101.5 | 101.0 |
| *hsp90ba* | 100.0 | 96.8 | 94.8 | 100.9 | 100.8 |
| *g6pd* | 97.0 | 92.0 | 91.8 | 97.8 | 98.1 |
| *ef1a* | 106.5 | 94.8 | 94.3 | 106.6 | 108.5 |
| *rpl13a* | 102.5 | 96.3 | 95.8 | 101.2 | 103.5 |

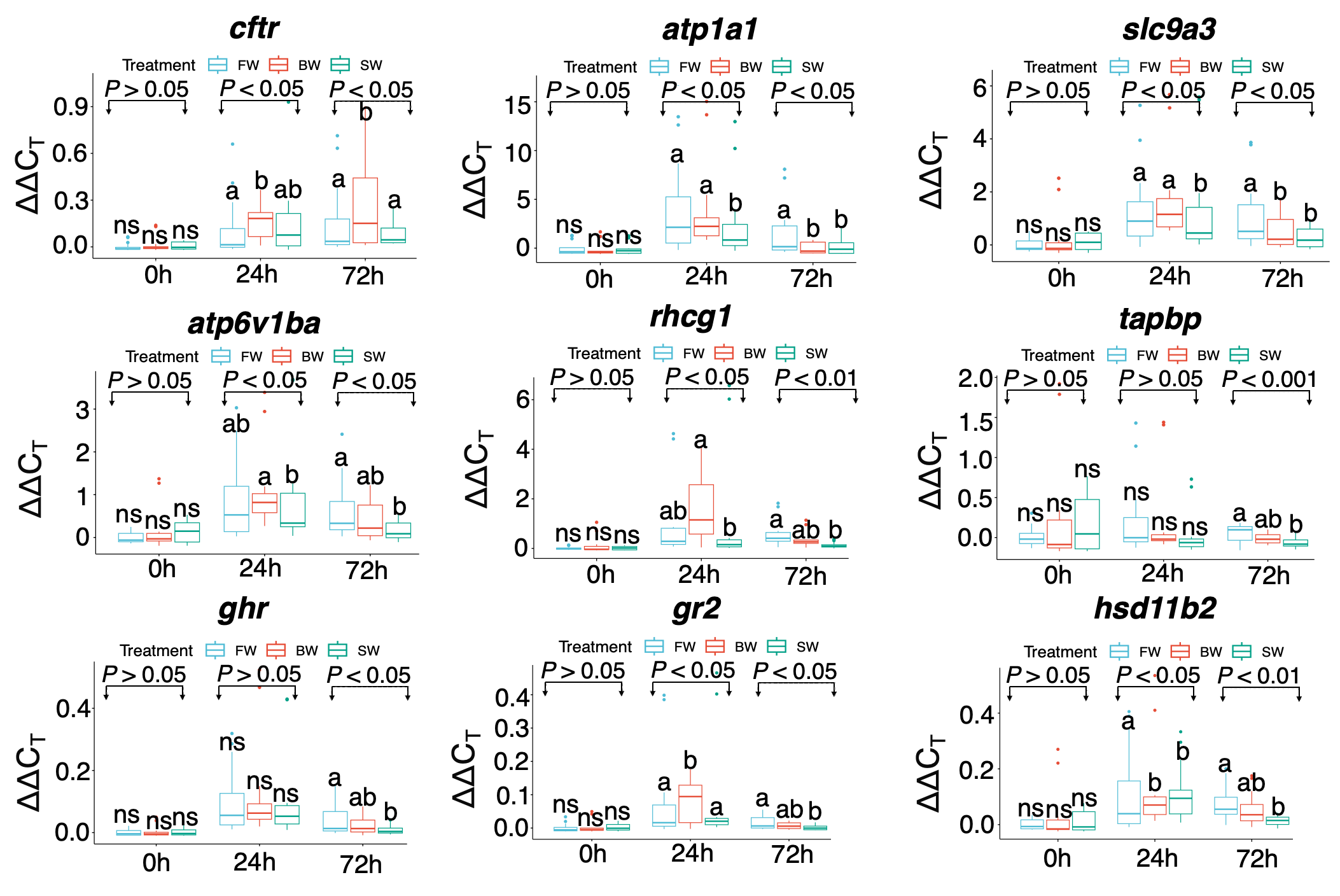

Supplementary Figure S1: Box plots represent nine genes that showed significant differences in transcript abundances in gill biopsy tissue based on mixed-effects models. The bold line represents the median, the boxes show the 25^th^ and 75^th^ percentiles, the whiskers indicate the 95% confidence interval, and the dots are outliers. Different letters denote significant differences in mean ΔΔC_T_ values among treatments.

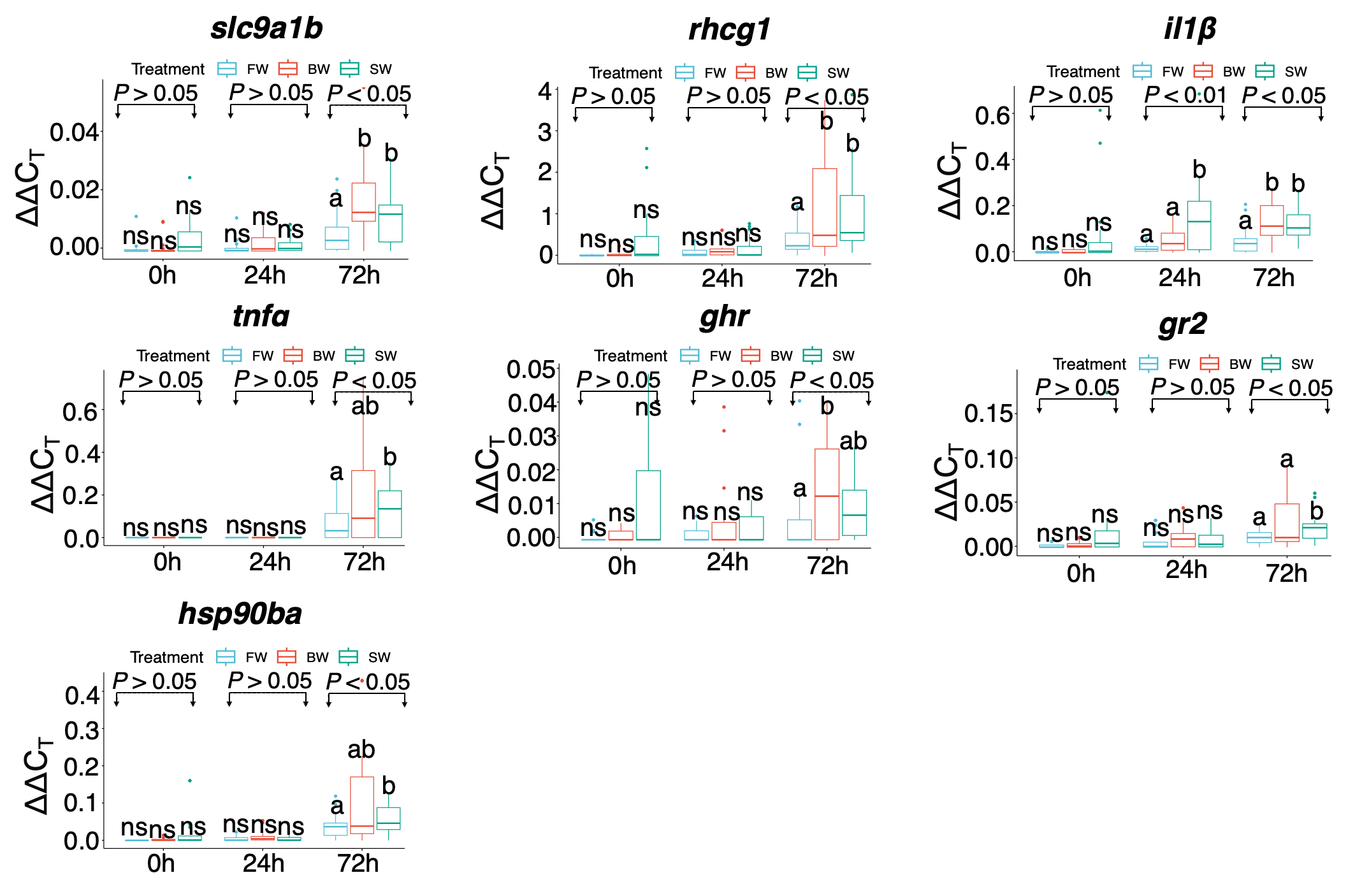
Supplementary Figure S2: Box plots represent seven genes that showed significant differences in transcript abundances in skin swab samples based on mixed-effects models. The bold line represents the median, the boxes show the 25^th^ and 75^th^ percentiles, the whiskers indicate the 95% confidence interval, and the dots are outliers. Different letters denote significant differences in mean ΔΔC_T_ values among treatments.

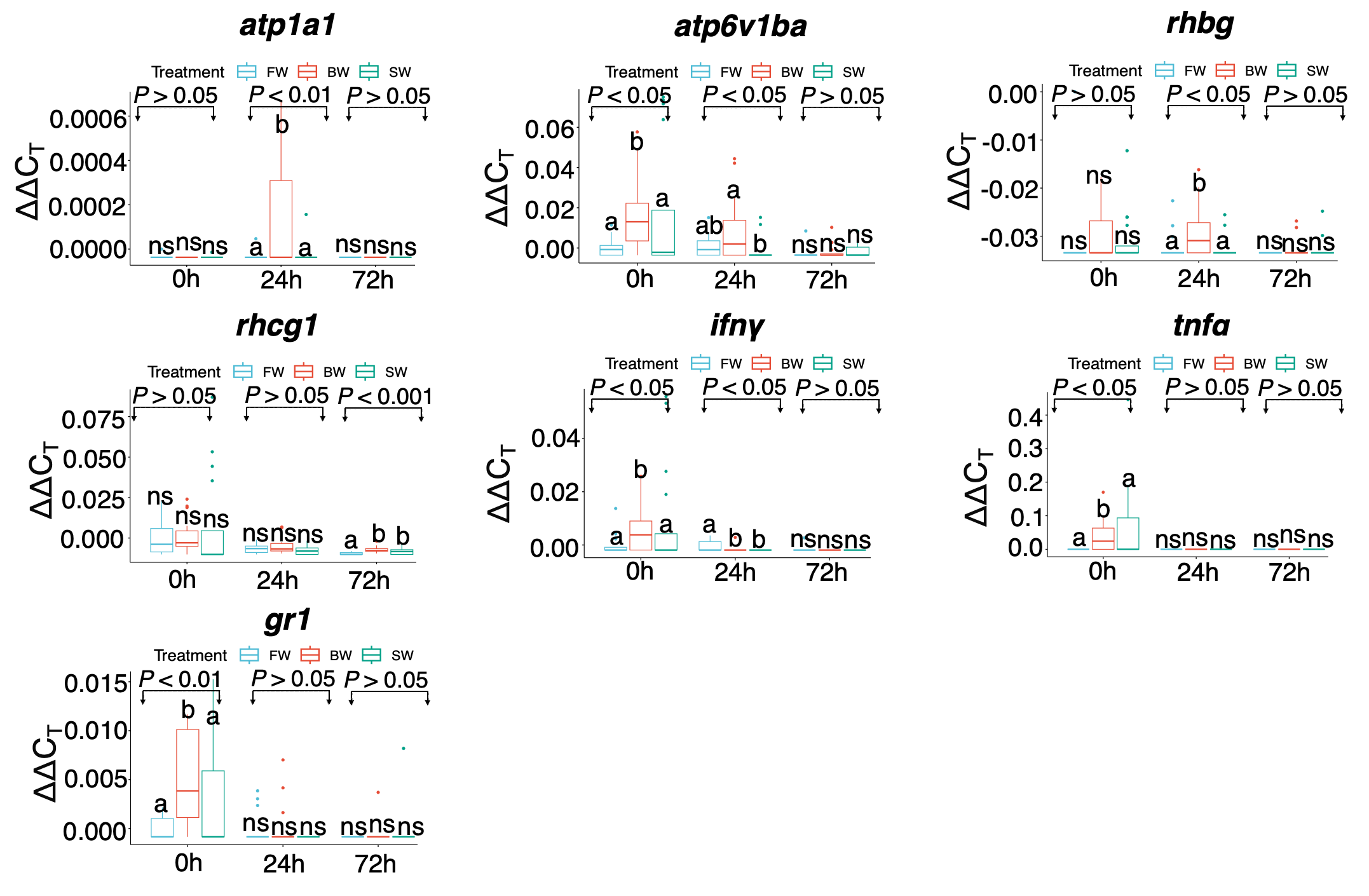
Supplementary Figure S3: Box plots represent seven genes that showed significant differences in transcript abundances in water (eRNA) samples based on mixed-effects models. The bold line represents the median, the boxes show the 25^th^ and 75^th^ percentiles, the whiskers indicate the 95% confidence interval, and the dots are outliers. Different letters denote significant differences in mean ΔΔC_T_ values among treatments.

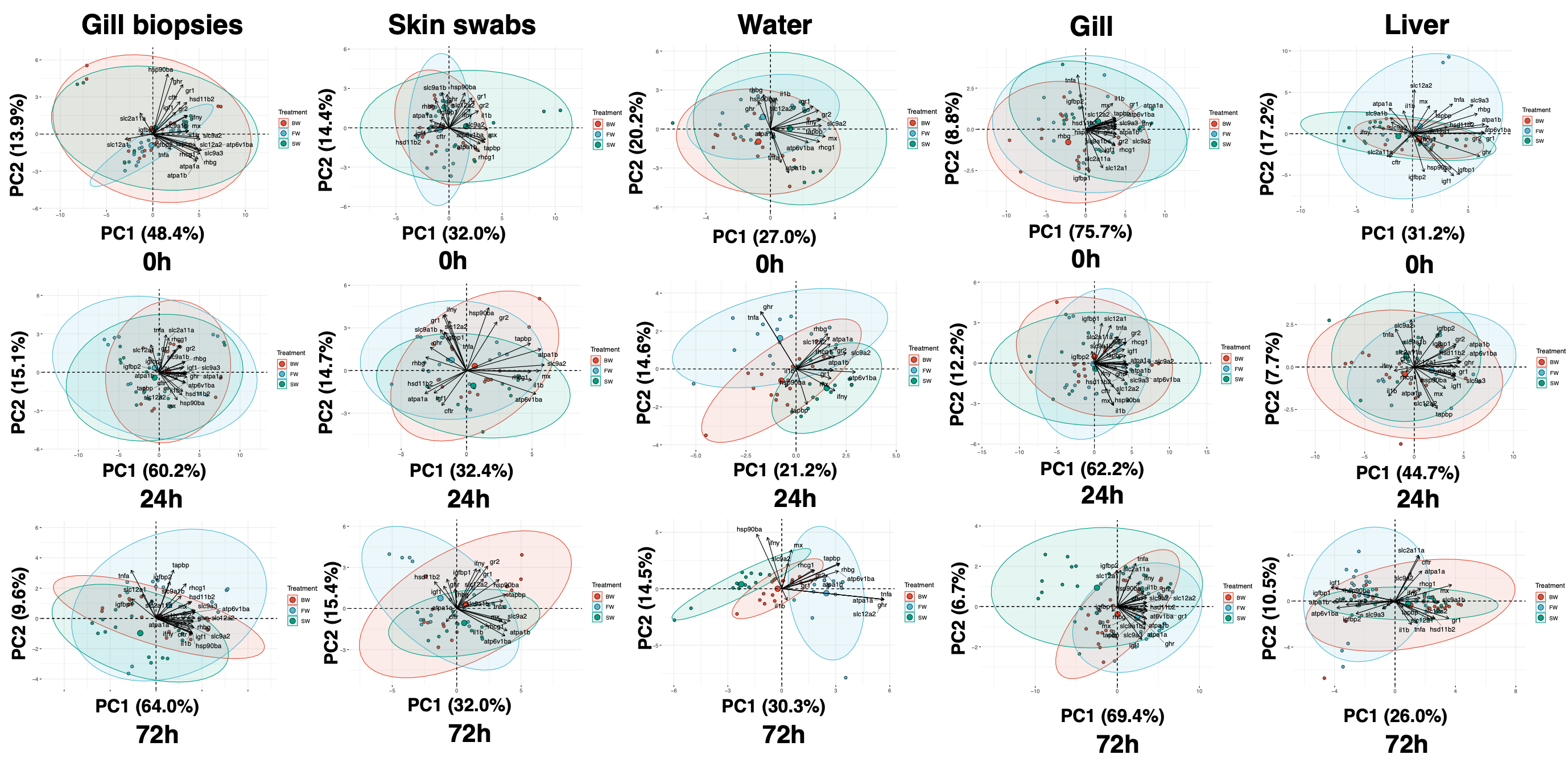

Supplementary Figure S4: Principal component analyses (PCA) of the qPCR assays that showed the mRNA transcript abundance across three time points (0 h, 24 h, and 72 h) across tissues (gill biopsies, skin swabs, water, gill, and liver). PCA plot illustrating the distribution of all three treatments (FW, BW, and SW) included in the study in the multivariate space. Loading vector plot showing the association of each candidate gene of interest with the variance explained by PC1 and PC2. Of the selected 25 assays, the assays that did not show amplifications either any of the five tissues for any treatment or time point were not included in the PCA.
